# Interactive analysis of Long-read RNA isoforms with Iso-Seq Browser

**DOI:** 10.1101/102905

**Authors:** Jingyuan Hu, Prech Uapinyoying, Jeremy Goecks

## Abstract

**Background:** Long-read RNA sequencing, such as Pacific Biosciences’ Iso-Seq method, enables generation of sequencing reads that are 10 kilobases or even longer. These reads are ideal for discovering splice junctions and resolving full-length gene transcripts without time-consuming and error-prone techniques such as transcript assembly and junction inference.

**Results:** Iso-Seq Browser is a Web-based visual analytics tool for long-read RNA sequencing data produced by Pacific Biosciences’ isoform sequencing (Iso-Seq) techniques. Key features of the Iso-Seq Browser are: 1) an exon-only web-based interface with zooming and exon highlighting for exploring reference gene transcripts and novel gene isoforms, 2) automated grouping of transcripts and isoforms by similarity, 3) many customization features for data exploration and creating publication ready figures, and 4) exporting selected isoforms into fasta files for further analysis. Iso-Seq Browser is written in Python using several scientific libraries. The application and analyses described in this paper are freely available to both academic and commercial users at https://github.com/goeckslab/isoseq-browser

**Conclusions:** Iso-Seq Browser enables interactive genome-wide visual analysis of long RNA sequence reads. Through visualization, highlighting, clustering, and filtering of gene isoforms, ISB makes it simple to identify novel isoforms and novel isoform features such as exons, introns and untranslated regions.

## Background

Isoform sequencing (Iso-Seq) is a long-read RNA sequencing method developed by Pacific Biosciences for finding and characterizing alternatively spliced gene isoforms [1]. Iso-Seq can generate long sequence reads more than 10 kilobases in length that may span from the isoform’s 5’ transcription start site all the way to its poly-A tail. This allows for direct isoform analysis without the need for isoform assembly and prediction, both of which have limited accuracy due to difficulties in combining short reads into complete isoforms [2,3].

Visual analysis of long isoforms is a challenge that existing bioinformatics tools do not address well. In most genome browsers, a large amount of space is taken up by intronic regions, making it difficult to view a complete isoform in detail and assess the presence or absence of exons and different splicing locations. To the best of our knowledge, there are only two tools devoted to visualization long-read RNA-seq data, MatchAnnot [4] and Iso-View [5]. MatchAnnot and IsoView solve this problem by removing the introns from view and vertically aligning matching exons making alternate isoforms easier to compare. In addition to visualization, automatically clustering isoforms based on similarity has proven useful for comparing isoforms, and Iso-View performs isoform clustering based on similar splicing patterns. However, these tools generate static images and lack dynamic features for interactively analyzing Iso-Seq data, such as exploring different genes, experimenting with different numbers of clusters, and saving selected data for additional analysis.

In this application note, we introduce Iso-Seq Browser (ISB), an interactive visual analytics tool for Iso-Seq data. ISB provides an exon-only Web-based interface for interactively viewing and clustering long-read RNA sequencing data. ISB draws several important features from MatchAnnot and IsoView and implements them in a dynamic, Web-based framework and also introduces new features as well. Key features of ISB include zooming and exon highlighting for comparing novel gene isoforms and reference transcripts, dynamic clustering of isoforms with transcripts, isoform filtering based on read support, creating publication-ready figures, and exporting data for further analysis.

## Implementation

There are two inputs for the Iso-Seq Browser (ISB): (1) annotated long RNA sequence reads and (2) a gene annotation. Annotated sequence reads are obtained by using MatchAnnot to combine a SAM file of aligned reads with gene annotation information to produce collections of reads that align to reference isoforms found in the annotation. ISB visualizes a single gene with all its reference and sequenced isoforms using an exon-only view. When a new gene is entered or selected, a new visualization is created. MatchAnnot assigns reads/isoforms to transcripts in a gene annotation using base-pair matching. This is often suboptimal because splicing patterns can be more important than base-pair matching. For this reason, ISB clusters isoforms and transcripts with k-means clustering using exon/intron locations, which better represent splicing patterns. Reference transcripts are black and isoform clusters are colored. A user can select the number of clusters, and ISB dynamically updates.

ISB provides several controls for exploring and refining visualized isoforms. Mousing over the visualization highlights exons and provides information about them. Isoforms can be filtered based on the number of full-length and partial length reads, making it simple to find isoforms with substantial supporting evidence and hide isoforms with limited support. Visualization width can be set directly, and overall visualization height is controlled by isoform/transcript height. Genes can be saved to easily return to them for further investigation, and isoform sequence data can be exported in fasta format for additional analysis.

ISB is written in Python2 using several Python scientific libraries, including Pandas, Scikit-learn, and the Bokeh visualization library. ISB is a Bokeh server application: the user interface, including interactio widgets and visualization, are viewed in a Web browser, but all data analysis is done on the server. This approach ensures maximum scalability and enables ISB to visualize and analyze datasets from entire sequencing runs and genome-wide annotations.

## Results and Discussion

The first step to use Iso-Seq Browser is to enter the name of either gene annotation file, or the annotated MatchAnnot reads files (which are in Python pickle format). The visualization begins when a specific gene is entered. The first plot will takes ~30-150 seconds to generate for large datasets because the gene annotations and annotated reads are loaded into server memory, but visualizations for subsequent genes are very fast. Genes with 40 or less isoforms can be visualized in ~1-3 seconds; genes with more than 40 isoforms will take longer to visualize due to additional time needed for clustering. After loading, the visualization is displayed along with a table of genes and their isoform counts. This table can be sorted to find genes with a large number of isoforms.

Figure 1 shows the visualization of the *BRCA1* (breast cancer 1) gene in the MCF-7 breast cancer cell line dataset available from Pacific Biosciences (http://www.pacb.com/blog/data-release-human-mcf-7-transcriptome/). *BRCA1* is a well-known gene widely implicated in breast cancer [6]. In the figure, black reference transcripts are interspersed with colored isoforms found in the MCF-7 Iso-Seq data. Eleven novel isoforms are clustered into 4 groups. To find isoforms with substantial supporting evidence, the slider for full reads support threshold is adjusted to 5. This filtering leaves only 2 isoforms that have 5 or more full-length reads that represent the entire isoform.

**Figure 1.**
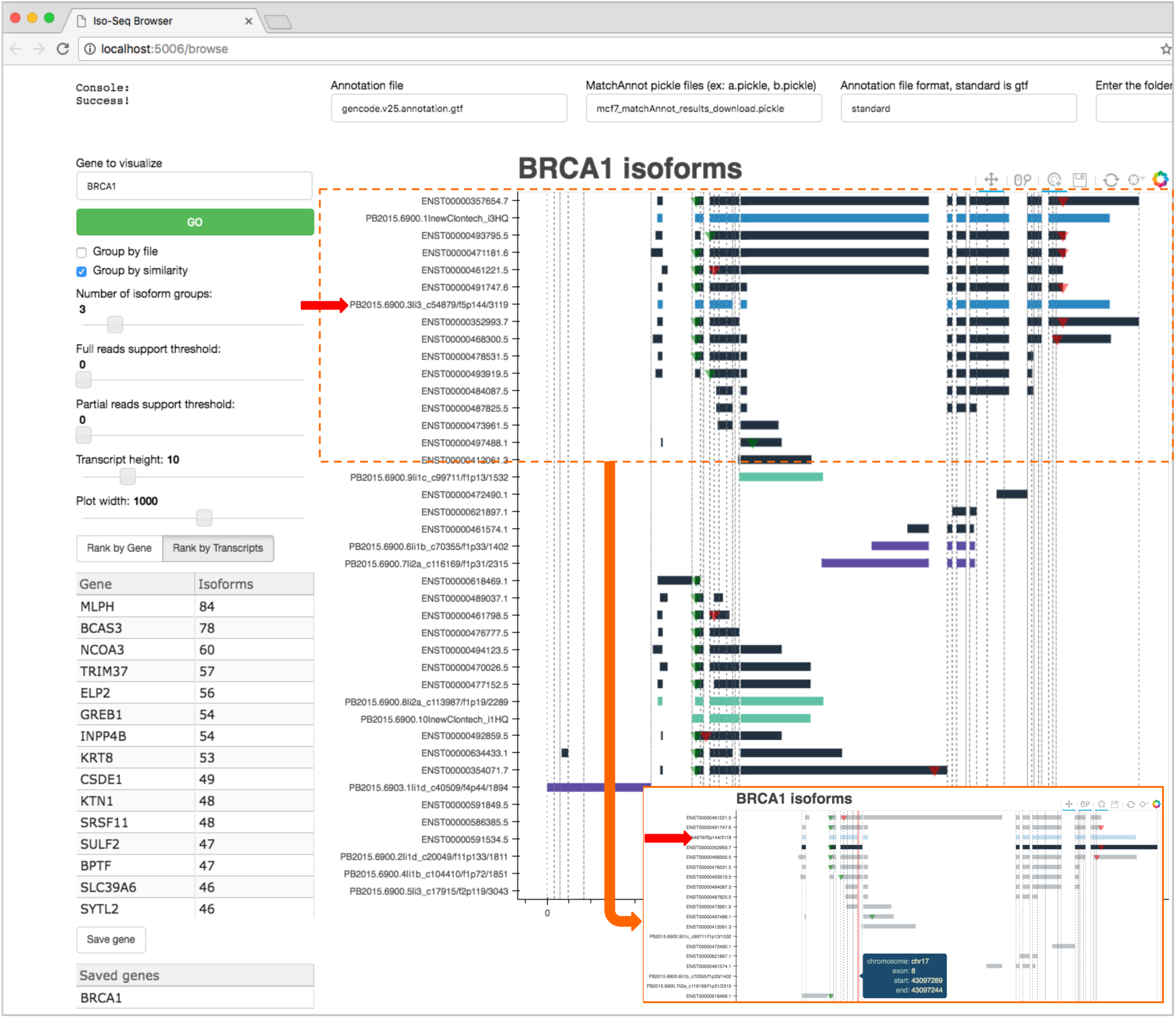
Visual analysis of long RNA-seq reads with the Iso-Seq Browser. BRCA1 gene isoforms and reference transcripts are visualized, and controls on the top and left provide options for showing, selecting, configuring, filtering, and saving isoform data. Inset: Full reads support threshold slider is used to show only isoforms comprised of 5 or more full reads, zoom is used to inspect an isoform (red arrow), and the tap tool is used to highlight a reference transcript and compare it to the isoform. Zooming and comparing to transcripts makes it clear that this is a novel isoform.

Zooming in on isoform PB2015.6900.3li3_c54879/f5p144/3119 near the top of the display, it appears that transcript ENST00000352993.7 below it is most similar to the isoform. Clicking on this transcript activates the tap tool, which shows transcript-specific exon and intron boundaries and numbering based on the selected transcript. Using the information from the tap tool, it’s clear that the isoform is missing exons 8 and 9 compared to the reference transcript. At any time, we can also save the current visualization view into a high-quality image file. In summary, we used ISB to visualize, filter, zoom, and compare sequenced isoforms with reference transcripts to easily identify a novel transcript and characterize its differences with the reference gene annotation.

## Conclusions

Iso-Seq Browser enables interactive Web-based, genome-wide visual analysis of long RNA sequence reads. Through visualization, highlighting, clustering, and filtering of gene isoforms, ISB makes it simple to identify novel isoforms and novel isoform features such as exons, introns and untranslated regions.

## Declarations

### Ethics

Not applicable.

### Consent to Publish

Not applicable.

### Competing Interests

The authors declare that they have no competing interests.

### Author Contributions

JH, PU, and JG conceived the project. JH and JG implemented the Iso-Seq Browser; PU and JG performed the data analysis discussed in the example vignette. JH, PU, and JG wrote the manuscript.

### Availability of data and materials

Project name: Iso-Seq Browser

Project home page: https://github.com/goeckslab/isoseq-browser

Code home page: https://github.com/goeckslab/isoseq-browser

Operating system(s): UNIX (Solaris recommended), Linux (Ubuntu or Debian recommended), MacOS (10.6+ recommended)

Programming language: Python

License: Academic Free

Any restrictions to use by non-academics: None

## List of Abbreviations

Iso-Seq: Pacific Biosciences’ isoform sequencing technology
ISB: Iso-Seq Browser
BRCA1: breast cancer 1 gene

## Acknowledgements

We thank Hiroki Morizono for his suggestions about how to improve the Iso-Seq Browser implementation.

